# HOXB6 and HOXB8 control immune-cancer cell interactions in pancreatic cancer

**DOI:** 10.1101/2024.09.06.611619

**Authors:** Ludivine Bertonnier-Brouty, Kavya Achanta, Jonas Andersson, Sara Bsharat, Tania Singh, Tuomas Kaprio, Jaana Hagström, Caj Haglund, Hanna Seppänen, Rashmi B Prasad, Isabella Artner

## Abstract

Pancreatic ductal adenocarcinoma (PDAC) is a lethal cancer lacking effective drugs and therefore new treatment targets are needed. Transcriptomic analysis comparing human embryonic and PDAC tissue identified a large overlap of expression profiles suggesting a re-initiation of developmental programs in pancreatic cancer. Specifically, we identified the transcription factors HOXB6 and HOXB8 as potential key regulators in PDAC. Loss of HOXB6 and HOXB8 in pancreatic cancer cells inhibited cell proliferation, induced apoptosis and senescence and enhanced gemcitabine sensitivity. Moreover, reduced HOXB6 and HOXB8 expression in pancreatic and lung adenocarcinoma cell lines affected transcription of immune response pathways which resulted in an increased sensitivity of cancer cells to anti-tumorigenic activities of macrophages suggesting that the HOXB6 and HOXB8 immune regulatory pattern is conserved in different cancer types. Additionally, naïve M0 macrophages exposed to HOXB8 deficient PDAC cells were unable to differentiate into tumor associated macrophages, suggesting that HOXB8 promotes the transition of initial anti-tumor macrophage to a tumor-promoting macrophage phenotype in pancreatic cancer. Our findings indicate that HOXB6 and HOXB8 play important roles in regulating cell proliferation, immune response and treatment resistance to promote pancreatic cancer tumorigenesis and could be useful therapeutic targets.

## INTRODUCTION

Pancreatic ductal adenocarcinoma (PDAC) is the most prevalent pancreatic cancer (1) with poor prognosis and low survival rate (< 10% 5-year survival rate). PDAC incidences have been on the rise, and the number of cases is projected to increase by 2-fold in the next decade (2–4). Low survival rates are exacerbated by the lack of effective treatment options. Current therapies often induce drug resistance, in part due to a dense tumor stroma, its specific immune cell microenvironment, and the presence of therapy-resistant cancer stem cells (5, 6). Thus, it becomes a priority to investigate the genetic and biological features of PDAC to facilitate diagnosis and the development of novel treatments.

Previous studies have defined molecular PDAC subtypes associated with different survival outcomes and involving differential expression of gene regulatory networks, which ultimately affect treatment sensitivity (7). According to these expression profiles and histopathological characteristics four PDAC subtypes were defined: squamous tumors mainly enriched for TP53/TP63/TGF-β signaling; pancreatic progenitor tumors preferentially expressing genes involved in early pancreatic development; immunogenic tumors with up-regulated expression of immune network genes; and aberrantly differentiated endocrine exocrine (ADEX) tumors which had increased expression of genes activating KRAS, as well as exocrine and endocrine differentiation.

The presence of a pancreatic progenitor PDAC subtype suggests that PDAC development and embryonic pancreas development share molecular similarities. Developmental regulators prevent apoptosis and ensure cell proliferation of progenitor cells. However, when these regulators are mis-expressed in adult cells, they can alter cell plasticity, promote the generation and survival of tumor cells thereby contributing to tumor formation (8). In line with this, elevated expression of transcription factors critical for pancreas development has been identified in the PDAC progenitor subtype. For example, SOX9 regulates pancreatic epithelial cell formation during embryogenesis (9) and is critical for maintaining ductal cell identity in the adult pancreas (reviewed in (10)). However, aberrant SOX9 expression has been detected in the majority of PDAC cases with poor prognosis and it has been shown that SOX9 expression promotes PDAC tumor formation (11). Interestingly, SOX9 regulates the expression of several pancreatic transcription factors critical for PDAC cells (GATA4, GATA6, FOXA2, HNF1A), further supporting the notion that transcriptional networks important for pancreas development are reinitiated. Most of the evidence provided above has been generated in animal models. However, human and mouse pancreas development differ significantly, restricting the possibilities of identifying common molecular pathways between human pancreas development and tumor formation. To identify novel signaling pathways in PDAC, a comparison of RNA sequencing data from human embryonic and adult pancreas with PDAC samples was performed, and expression profiles in specific PDAC subtypes were assessed. This analysis uncovered a significant enrichment of HOX transcription factors in PDAC. HOX genes are critical for embryogenesis, particularly for anterior-posterior patterning, but are also instrumental in tumor formation (12). Here, we show that the transcriptional regulators HOXB6 and HOXB8 are enriched in the embryonic and PDAC transcriptome with predominant expression in embryonic mesenchyme and malignant ductal cells. Loss of HOXB6 and HOXB8 in PDAC cells resulted in impaired proliferation and colony formation, increased cell viability, accompanied by increased apoptosis and senescence, as well as increased sensitivity to gemcitabine. Expression of gene networks regulating these processes was also dysregulated in HOXB6 and HOXB8 knocked down PDAC cells. Moreover, aberrant expression of genes influencing tumor-immune cell interactions was observed. Co-culture of HOXB6 and HOXB8 deficient PDAC cells with macrophages showed that HOXB6 and HOXB8 promoted tumorigenic properties of tumor-associated macrophages and protected PDAC cells from macrophage detection. Changes in immune evasion properties were also observed in a lung adenocarcinoma cell line, describing a novel function of HOX genes in tumor cell propagation and immune evasiveness.

## MATERIALS & METHODS

### Gene expression profiling in human fetal tissues and comparison with adult pancreas

Embryonic pancreata were obtained from terminated fetuses (7–14 gestational weeks, n=16). Informed consent was obtained from the participating women. Ethical permission has been obtained from the regional ethics committee in Lund (Dnr2012/593, Dnr 2015/241, Dnr 2018-579).

Bulk RNA was extracted for RNA-Seq, libraries generated, and comparison of fetal-specific gene expression with adult pancreas expression were performed. For comparison with adult tissues, data from GTEX was used. Raw data (bam files) were downloaded from the GTEX database for comparison with fetal datasets. To ensure consistency in data processing between fetal and adult samples, the RNA-Seq expression quantification pipeline from GTEX V8 (https://github.com/broadinstitute/gtex-pipeline/) was used on both datasets. Paired-end 101 bp-long reads were aligned to the Reference Human Genome Build 38 using STAR v2.5.3a, to the reference genome annotation Gencode v26. After alignment and post-processing, expression quantification was performed following the pipeline using RSEM and RNASeqQC. This resulted in count matrix normalized for library sizes. Normalization of library sizes was performed in edgeR by dividing counts by the library counts sum and multiplying the results by a million to obtain counts per million (CPM) values.

Comparison with expression in adult pancreas: edge-R was used to perform differential expression analysis with age and sex as covariates for the genes of interest. Batch correction was performed by considering platform and batches using COMBAT.

Single-cell RNAseq of embryonic pancreas were performed and analyzed as previously described (13, 14).

### Gene expression studies in PDAC samples

Sequencing data consisting of PDAC samples (n=24) and control samples (n=11) from Peng et al (15) were used (accession number: CRA001160) and processed as closely as possible to the parameters described previously (15). Ten cell types were identified and categorized among the PDAC samples separately according to the criteria used by Peng et al (15). These cells were re-clustered in Loupe browser with default settings to get a tSNE plot, which resulted in a total of 57383 re-clustered cells (PDAC= 41862 and control=15521). HOX expression was then filtered for, and plots were generated as described previously (14).

### Cells

The human pancreatic cancer cell line PANC-1 (16) and the human monocytic leukemia cell line THP-1 (17) were obtained from the European Collection of Authenticated Cell Cultures (Sigma-Aldrich, MO, USA, 87092802 and 88081201). The lung adenocarcinoma cell line Calu-3 was a gift from Darcy Wagner, Lung Bioengineering and Regeneration group, Lund University. PANC-1 cells were maintained in DMEM high glucose medium (Sigma-Aldrich, D6429) supplemented with 10% FBS and 1% penicillin–streptomycin (Gibco, 15140122) and THP-1 cells in RPMI-1640 medium (Gibco, 11875-093) supplemented with 10% FBS and 1% penicillin–streptomycin-neomycin (Gibco, 15640055). Calu-3 cells were maintained in DMEM low glucose medium (Gibco, 31885023) supplemented with 10% FBS and 1% penicillin–streptomycin (Gibco, 15140122). Cells were incubated at 37°C in a humidified incubator with 5% CO_2_ and frequently tested for mycoplasma contamination using MycoAlert™ detection kit (Lonza, LT07-118).

PANC-1 and Calu-3 cells were transfected with Silencer™ Select negative control siRNA (Invitrogen, USA, 4390843), Silencer™ Select HOXB6 siRNA (Invitrogen, s6805) and/or Silencer™ Select HOXB8 siRNA (Invitrogen, s6810) at 10 nM with RNAiMax Lipofectamine (Invitrogen, 13778150) according to the manufacturer’s guidelines for 48h or 7 days. Transfection efficiency was validated using Silencer™ Cy™3-labeled GAPDH siRNA (Invitrogen, AM4649).

### RNA-seq of PANC-1 cells

Seven days after transfection, RNA was extracted, RNA libraries constructed, and RNA sequencing performed on a NovaSeq 6000 System (Illumina, USA). Data quality assessment was done using FastQC tool and the quality control analysis was performed using the Cutadapt tool (version 4.4). STAR (Spliced Transcripts Alignment to a Reference) tool was used to align reads with human reference genome (hg38). Count data of expressed genes were generated using hg38 annotation gtf file and then subjected to differential gene expression analysis using R package DESeq2 (19). The filtering criteria used for obtaining list of statistically significant differentially expressed genes was p-value < 0.05 and Log2FC > 1 or < -1 (for upregulated or downregulated genes, respectively). Single Sample Gene set Enrichment Analysis (ssGSEA) method is an extension of the GSEA method, working at the level of a single sample rather than a sample population as in the original GSEA application. The score derived from ssGSEA reflects the degree to which the input gene signature is coordinately up- or downregulated within a sample.

### TIMER

Correlations between immune infiltration level and HOXB6 or HOXB8 expression in pancreatic and lung adenocarcinoma were obtained from TIMER 2.0 immune association module (18) (http://timer.cistrome.org/).

### Study population and tumor tissue microarray analyses

We studied a cohort of 154 patients with pancreatic adenocarcinoma described previously (14). The Surgical Ethics Committee of the Helsinki University Hospital (HUH) approved the study protocol (Dnr. HUH 226/E6/06, extension TKM02 §66 17.4.2013).

Immunohistochemistry of paraffin sections was performed using HOXB6 (219499, Abcam, England, 1:200) or HOXB8 (bs6339R, Thermo Fischer, USA, 1:200) antibodies. Immunoreactivity in tumor cells was interpreted independently by two researchers (T.K. and J.H), while the researchers were blinded to the clinical outcome of the patients.

Negative staining was scored as 0, weakly positive as 1, moderately positive as 2, and strongly positive as 3 separately for tumor cells for both HOXB6 and HOXB8 and overall disease-specific survival (DSS) analyses were performed as described previously (14).

### qPCR

Total RNA was extracted using RNeasy Qiagen kit and cDNA was generated. qPCR assays were performed using StepOnePlus Real-Time PCR System. Relative gene abundance was calculated using the ΔΔCt method and expressed as FC to control (Table S1), and used TBP and ribosomal gene S18 as a housekeeping gene.

### Colony assay

Colony formation assay was performed as previously described (14) and analyzed by ColonyArea plugin (19) with ImageJ software. Colony formation was quantified by determining the percentage of area covered by cell colonies and the cell density according to the intensity of the staining.

### Proliferation

Proliferation was quantified using Click-iT™ Plus EdU Cell Proliferation Kit for Imaging (Invitrogen, C10640). Cells were prepared as described previously (14) and EdU labelling was done according to manufactureŕs instructions (Invitrogen, C10640).

### Triplex assay

PANC-1 cells were prepared as described previously (14) and ApoTox-Glo triplex assay kit (Promega, WI, USA, G6320) was used according to manufacturer’s instructions.

### Senescence

Seven days after transfection in 8-well chamber slides, senescence-associated β-galactosidase staining was done following the Crowe et al (20) detailed protocol. Cells were fixed and incubated with freshly prepared SA β-gal stain containing X-gal (Roche, 11680293001) for 16h at 37°C in an incubator without CO_2_. Then cells were washed and stained with DAPI. Images were acquired using a slide-scanner microscope (Olympus VS120) and quantification was done using the Cell Counter Plugin in ImageJ to manually tag and count stained cells for each colour channel. Blue-colored cells were counted as senescent cells and a ratio was determined according to the total number of cells.

### Gemcitabine

Cells were transfected with siRNAs in 96 well plates at a density of 1×10^3^ cells/well (siCtrl), 1.5×10^3^ cells/well (siHOXB6 and siHOXB6B8) or 3×10^3^ cells/well (siHOXB8) to obtain the same cell density 7 days after transfection. 7 days after transfection, cells were exposed to increasing concentration of gemcitabine (Sigma-Aldrich, G6423) for 6h. Then, cells were washed and maintained in complete DMEM medium for 96h. Cell number was determined using the cell proliferation kit I (MTT, Sigma-Aldrich, 11465007001). The concentration of gemcitabine required to inhibit cell proliferation by 50% (IC50) was calculated.

### Generation of M0 and TAM macrophages from THP-1 cells and medium collection

THP-1 cells were treated with phorbol 12-myristate 13-acetate (PMA, Sigma-Aldrich, P8139) at 100 ng/ml for 24h in 6 well plates (1.5×10^6^ cells/well) or 6-well inserts (0.4µm pore size, Falcon, 353090, 6.5×10^5^ cells/insert). Activated THP-1 cells were washed and maintained in complete RPMI-1640 medium for another 24h. Cells were considered M0 macrophages at this stage. Medium was changed again and M0 conditioned medium were collected after 24h of incubation. PANC-1 or Calu-3 cells were transfected, incubated with M0 or control conditioned medium and cell number was determined (MTT kit, Sigma-Aldrich, 11465007001).

For the generation of specific-KD tumor-associated-macrophages (TAM), M0 macrophages were activated in 6-well inserts and then co-cultured with transfected PANC-1 or Calu-3 cells for 72h. Then, macrophages were isolated, medium was changed, and TAM conditioned medium was collected after 24h of incubation. PANC-1 or Calu-3 cells were incubated with associated TAM, M0 or control conditioned medium and cell number was determined (MTT kit, Sigma-Aldrich, 11465007001).

Collected media were centrifuged at 1200[rpm for 5 min to remove cell debris, filtered using 0.2μm syringe filters (Sarstedt, 83.1826.001), and stored at -80[°C. For experimental procedures, all conditioned media were diluted 1:1 with fresh media.

### Statistics

Statistical analyses and graphs were performed using R software with ggplot2 and ggpubr packages. Once normality and homoscedasticity of the variances verified, multiple comparisons between control and different siRNAs were analyzed using ANOVA with Tukey’s post-hoc test.

## RESULTS

### Identification of genes controlling human pancreas development and pancreatic cancer

Previous studies have shown that expression of developmental regulators is reinitiated in PDAC (8). RNA sequencing of 16 fetal pancreata (7-14 weeks post conception) was performed to identify novel genes critical for embryonic pancreas development. To identify key genes associated with PDAC, we selected two independent studies (21, 22) that reported gene expression signatures and pathways in this disease. Yan et al. identified 2735 differentially expressed genes in pancreatic tissue from cancer patients compared to adjacent benign tissue, with 557 of these genes upregulated. Similarly, Mao et al. found 2725 differentially expressed genes, of which 1554 were upregulated.

The upregulated genes from both studies were compared to determine which genes were robustly and reproducibly altered in PDAC. This resulted in 332 genes that were differentially expressed in both studies (Fig.1A).

**Figure 1.**
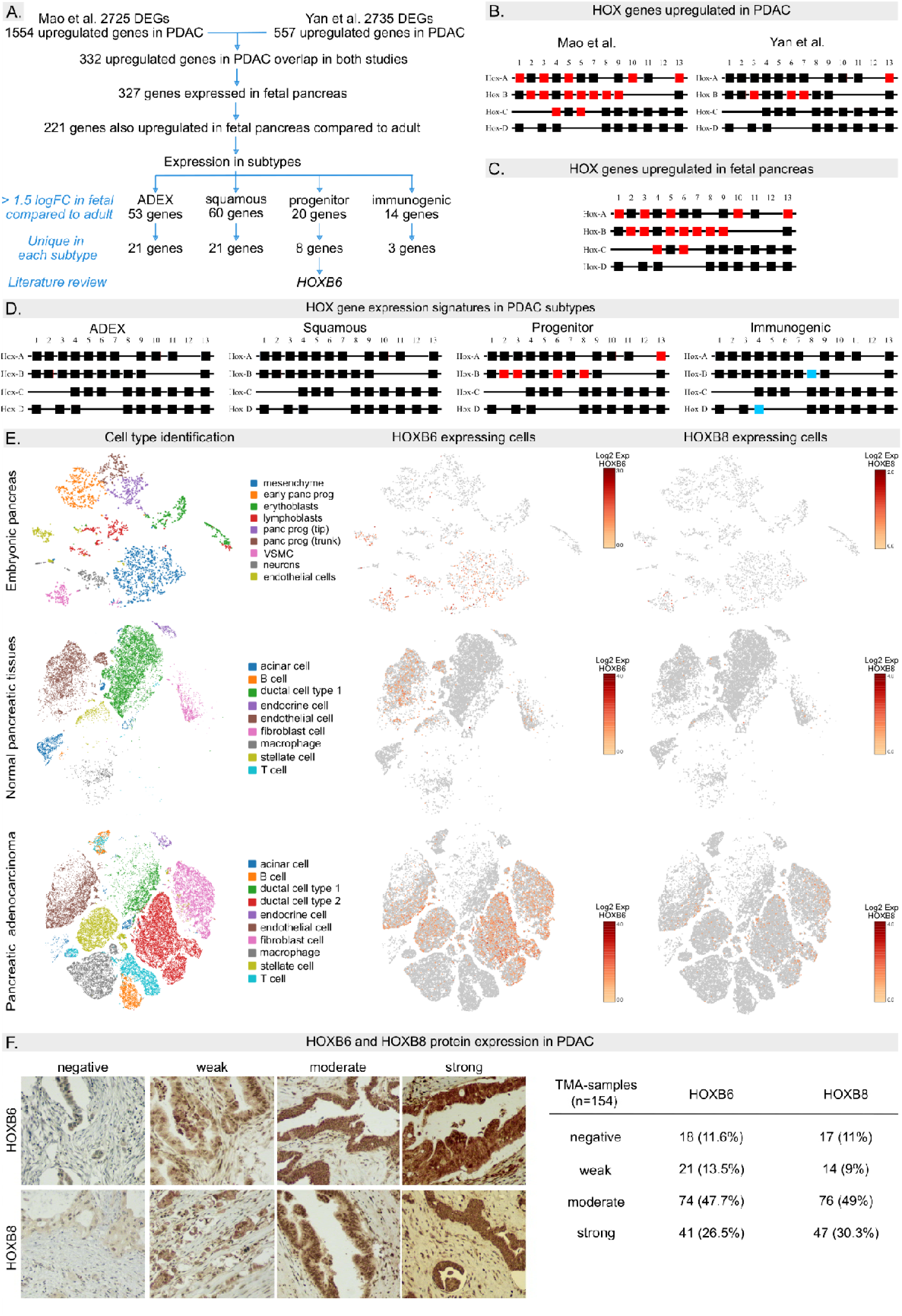
HOXB6 and HOXB8 are upregulated in fetal and PDAC tissue. (A) Flow chart illustrating the workflow for identification of genes of interest. (B) Squares represent HOX genes organized by clusters. HOX genes which are upregulated in PDAC in Mao et al. and Yan e al. studies are indicated in red. (C) HOX genes upregulating in fetal pancreas in red. (D) HOX gene expression signatures in PDAC subtypes. Genes upregulated or downregulated are indicated in red and blue, respectively (E) t-SNE embedding and HOX gene expression in pancreas from an 8-week PC embryo, in normal adult pancreas and PDAC tissues. Cells with HOXB6 or HOXB8 expression were colored according to their expression levels. (F) Representative images of HOXB6 and HOXB8 immunohistochemical staining and distribution of TMA-samples according to staining intensity. Original magnification: 200x.

To identify the intersect of genes which were upregulated in both pancreatic cancer and fetal tissue compared to adult, gene expression data from embryonic pancreas was first compared with expression data from adult pancreas obtained from GTEX to determine genes that were enriched in pancreas anlagen. 327 genes were expressed in the fetal pancreas (221 genes with elevated expression of which 69 were exclusively expressed in fetal pancreas).

Next, we assessed if genes enriched in embryonic pancreas and PDAC clustered in specific PDAC subtypes (7) with the majority of the genes belonging to squamous (60) and ADEX (53) subtypes (Fig.1A, listed in table S2). Many of these genes have been studied in the context of PDAC biology (table S3) confirming the validity of our approach. Several differentially expressed genes belong to the HOX family of transcription factors (Fig.1B) and have been studied in PDAC development (23–25). However, the function of HOXB6 (enriched in progenitor PDAC subtype, Fig.1B-D) and the closely related (26) HOXB8 transcription factor (differentially expressed in progenitor and immunogenic PDAC subtype) has not been investigated in PDAC so far.

### Localization of HOX candidate genes in developing pancreas and PDAC tissues

Previous studies have detected HOXB6 in mesenchymal cells during mouse pancreas development (27), with its expression restricted to fibroblasts in the adult pancreas (28). Meanwhile, no expression analysis of HOXB8 in the pancreas has been reported yet. We assessed the expression HOXB6 and HOXB8 in developing human pancreas and PDAC tissue using scRNAseq (13, 15). During pancreas development, HOXB6 was predominantly expressed in mesenchymal cells, whereas HOXB8 was only present in a few mesenchymal cells as well as in vascular smooth muscle cells (Fig.1E). In the adult pancreas, HOXB6 expression was detected mainly in fibroblasts, endothelial and stellate cell populations (3.8% of all pancreatic tissue cells), while HOXB8 expression was limited (0.18% of all pancreatic tissue cells). In contrast, HOXB6 transcripts were abundant in PDAC tissues (12% of all PDAC cells). mRNA expression was mainly detected in malignant ductal (23% ductal cell type 2), endothelial (20%) and fibroblast cells (16%). HOXB8 was expressed in the same cell clusters but in a lower proportion (2.8%, 2.5% and 3.5% HOXB8^+^ cells, respectively) (Fig.1E).

Tissue microarray analysis (TMA) was performed to confirm gene expression and to assess if HOXB6 and/or HOXB8 expression were correlated with clinical parameters and patient survival. HOXB6 and HOXB8 protein expression was detected in most TMA-samples (88.3 and 89% respectively, Fig.1F). Kaplan–Meier analysis revealed that low HOXB6 expression was associated with poorer disease specific survival compared to moderate expression or high expression (Fig.S1, table S4). These findings show that HOXB6 and HOXB8 proteins are expressed in the vast majority of all PDAC samples and that variations in protein expression are associated with altered survival rates.

To determine which gene networks were co-expressed with HOXB6 and/or HOXB8 in PDAC cells, an unbiased co-expression analysis of single fetal and normal adult cells was performed. HOXB6 and HOXB8 expression in fetal and PDAC tissue was correlated with genes regulating RNA and protein metabolism, DNA replication, signal transduction, programmed cell death, cell cycle regulation, and stem cell biology (table S5-8) suggesting that a large subset of HOXB6 and HOXB8 expressing cells are pancreatic cancer stem cells and that HOXB6 and/or HOXB8 regulate stemness, proliferation and apoptosis in PDAC cells.

### Loss of HOXB6 and HOXB8 expression affect clonogenic capacity, cell proliferation, and viability of PANC-1 cells

Co-expression analysis showed that HOXB6 and HOXB8 expression correlate with genes regulating cell cycle, proliferation, and apoptosis. To determine if these observations were functionally relevant in PDAC cells, we knocked down (KD) HOXB6 and/or HOXB8 expression using siRNAs (i.e., siHOXB6, siHOXB8 and siHOXB6B8) in the human pancreatic cancer cell line PANC-1. PANC-1 cells have been isolated from a pancreatic carcinoma of ductal cell origin and highly express HOX candidate genes. KD efficiency of siHOXB6, siHOXB8 and siHOXB6B8 was assessed by RT-qPCR (Fig.S2).

First, we examined if reduced HOXB6 and HOXB8 expression affected clonogenic capacity of tumor cells by performing colony formation assays. 7 days post transfection, the area and density of PANC-1 colonies were significantly decreased in all KD conditions compared to the control: in mean, colony formation was reduced by 35% in siHOXB6B8, 30% in siHOXB6 and 65% in siHOXB8 (Fig.2A).

**Figure 2.**
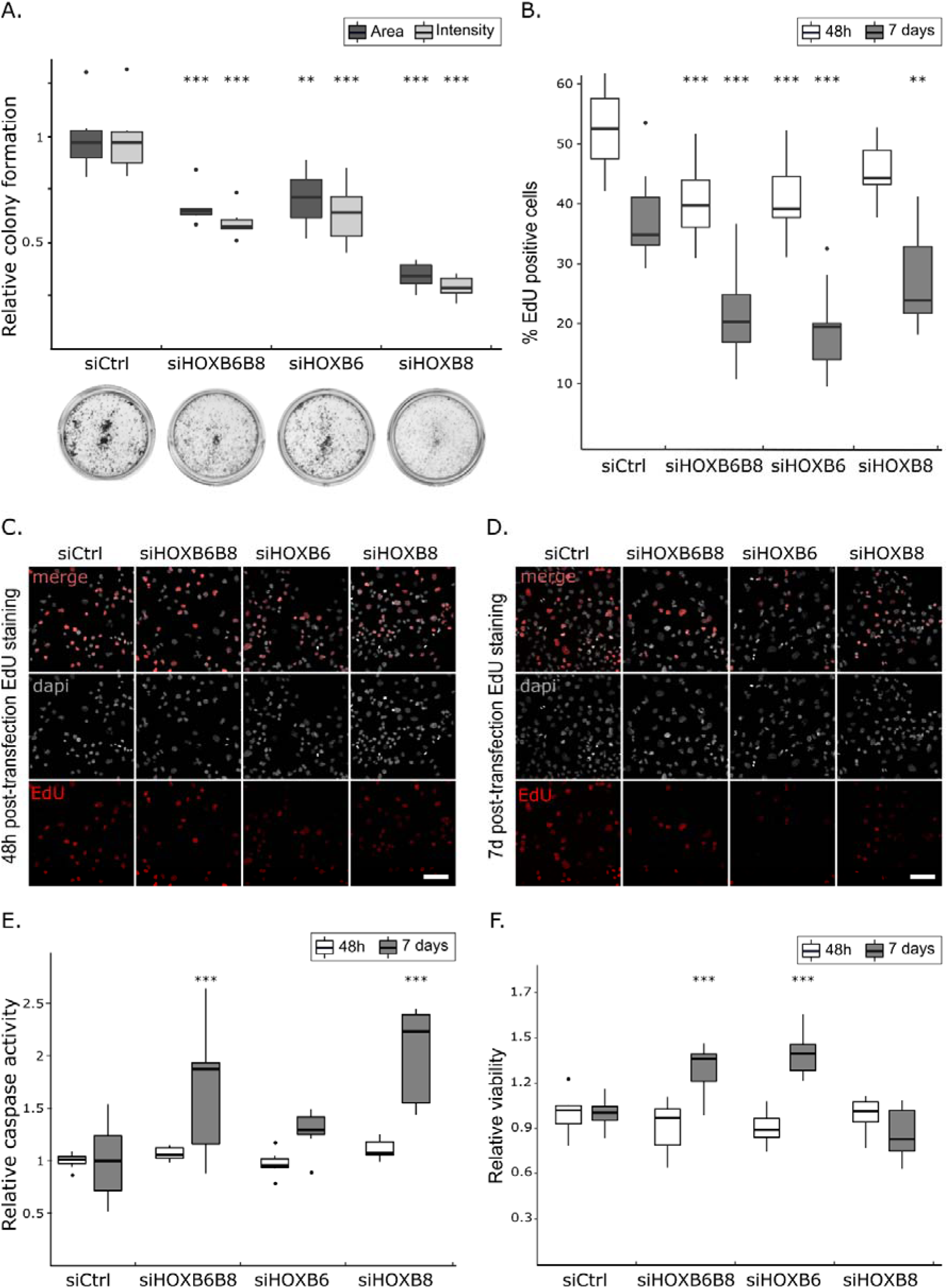
Colony formation, proliferation, cell viability and apoptotic capacities of siHOXB6 and SiHOXB8 transfected cells. Relative colony formation quantification (A). Results are shown as colony area or staining intensity percent 7 days post knock-down compared to negative control (scrambled RNA, siCtrl). (B) Quantification of EdU positive cells. Results are shown as percent EdU positive cells of total cell number. Dapi and EdU staining 48h (C) or 7 days (D) after transfection. Scale bars = 100 µm. Cells were exposed to EdU for 4h. (E) Cell viability 48h and 7 days post transfection. Results are shown as viability/cytotoxicity ratio compared to negative control. (F) Relative caspase activity 48h or 7 days after transfection. Results are shown as caspase-3/7 activity compared to negative control. (A-F) Measurements were derived from three independent experiments with three replicates each. (A-B, E-F) Tukey’s post-hoc test significances are indicated by stars compared to control when significant. * p < 0.05, ** p < 0.01 *** p < 0.001. The box extends from the first to the third quartile (Q1/Q3) with a line representing the median. The whiskers extending from both ends of the box indicate variability outside Q1 and Q3. Everything outside is represented as an outlier.

In order to assess if siHOXB6 and/or siHOXB8 impaired proliferation of tumor cells, EdU incorporation was assessed as a measure of DNA synthesis (S-phase). At 48h post transfection, the proportion of EdU^+^ cells was significantly decreased in siHOXB6B8 and siHOXB6 compared to control cells (Fig.2B, C). Cell proliferation capacities were also tested 7 days post transfection to assess long term effects of reduced HOXB6 and HOXB8 expression. EdU^+^ cell number was reduced by 41% in siHOXB6B8, 48% in siHOXB6 and 24% in siHOXB8 cells compared to control (Fig.2B, D).

Colony formation of siHOXB8 cells was significantly more reduced compared to siHOXB6 and siHOXB6B8. These differences in colony formation between KD conditions cannot solely be explained by differences in the proliferation rate. To better understand the different phenotypes in siHOXB6 and siHOXB8, apoptosis and cell viability were assessed using ApoTox-Glo triplex assay. At 48h post transfection, no significant difference in caspase activity and the viability rate was observed (Fig.2E, F). At 7 days post transfection, caspase activity was significantly higher in siHOXB6B8 (+65%) and siHOXB8 (+100%) cells compared to control (Fig.2E). The apoptosis rate was significantly increased in siHOXB8 compared to siHOXB6 cells suggesting that differences in apoptosis rates resulted in decreased colony formation of siHOXB8 cells.

siHOXB6B8 cells had lower proliferation abilities and similar apoptosis rate, but more colonies than siHOXB8 cells. To assess if these changes were associated with differences in cell viability, Triplex assays were performed 7 days post transfection. siHOXB6 (+39%) and siHOXB6B8 (+28%) cells showed significantly increased cell viability compared to control cells whereas cell viability wasńt affected in siHOXB8 (Fig.2F).

### Hox gene deficiency enhances senescence and gemcitabine sensitivity

Decreased proliferation rates and colony formation, but increased cell viability of siHOXB6 and siHOXB6B8 PANC-1 cancer cells may result from altered cellular senescence which is characterized by stable viability with resistance to apoptosis (29). siHOXB6 and siHOXB8 cells were assessed for galactosidase activity as a measure of cellular senescence. At 7 days post transfection, siHOXB6 (+107%), siHOXB8 (+137%) and siHOXB6B8 (+159%) cells displayed significantly higher galactosidase activity than control cells (Fig.3A, B). This demonstrates that HOX KD cells become senescent more readily.

**Figure 3.**
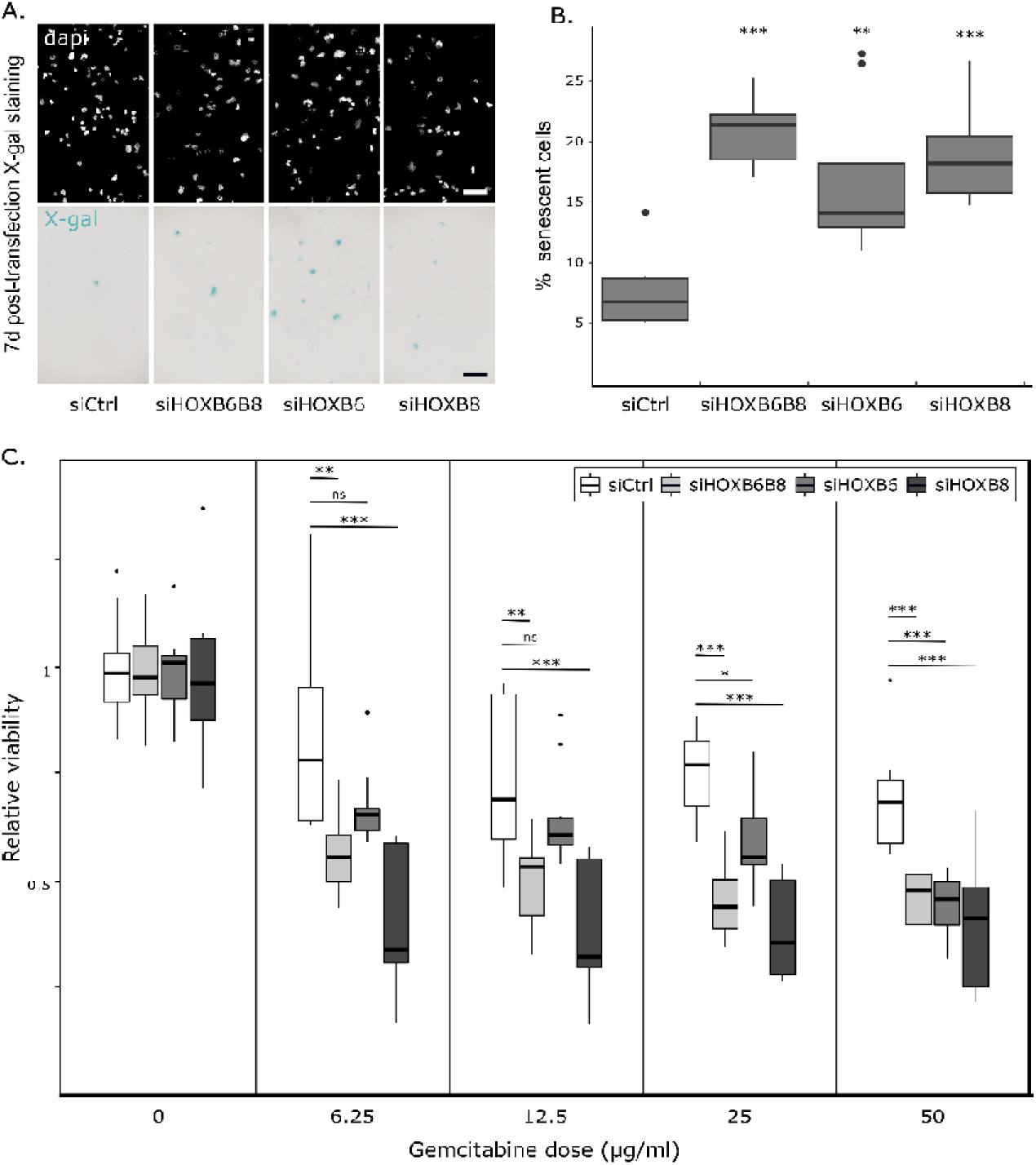
Senescence and gemcitabine sensitivity. (A) Senescence-associated galactosidase activity for siHOXB6, siHOXB8, and siHOXB6B8 is shown in blue. Scale bar = 100µm. B) Quantification of senescent cells for all conditions as percent per total cell number. (C) Relative viability of 7-days transfected cells treated with varying concentrations of gemcitabine. MTT assay quantifying the relative cell viability after 6h-treatment with varying concentration of gemcitabine (µg/ml) to the dose 0 for each transfected cell condition. Measurements were derived from three independent experiments with three replicates each. Tukey’s post-hoc test significances are indicated by stars compared to the control when significant. ns p > 0.05, * p < 0.05, ** p < 0.01 *** p < 0.001. The box extends from the first to the third quartile (Q1/Q3) with a line representing the median. The whiskers extending from both ends of the box indicate variability outside Q1 and Q3. Everything outside is represented as an outlier.

Senescence of cancer cells is associated with treatment resistance (30, 31). To determine if the increased cellular senescence of KD cells was linked to pancreatic cancer treatment resistance, we assessed sensitivity to gemcitabine, the primary pancreatic cancer treatment. In a dose-response experiment with Gemcitabine, siHOXB6 and/or siHOXB8 cells were more sensitive to treatment than control cells (Fig.3C). The concentrations of gemcitabine required to reduce cell number by 50% (IC50) were significantly lower for KD cells (IC50 siCtrl: 105ug/ml, IC50 siHOXB6: 40µg/ml (-61%), IC50 siHOXB6B8: 29µg/ml (-72%), IC50 siHOXB8 < 6.25 µg/ml (>-95%)) suggesting that HOXB6 and HOXB8 function in PDAC cells contributes to gemcitabine independently from its role in cellular senescence.

### HOX genes regulate cell cycle, proliferation and senescence pathways

HOXB6 and HOXB8 transcription factors have multiple roles in gene regulation (32). To assess how HOXB6 and/or HOXB8 transcriptional activity affects gene expression in pancreatic cancer cells, RNA sequencing of siHOXB6 and/or siHOXB8 PANC-1 cells was performed. Analysis of siHOXB6, siHOXB8 and siHOXB6B8 samples indicated altered expression for 5611, 5905 and 7384 genes, respectively (tables S15-17). 2553 common genes were dysregulated in all conditions.

To identify novel cellular processes and functions associated with HOXB6 or HOXB8 expression, differentially expressed genes were analyzed using MSigDB (33, 34), Gene set enrichment analyses (GSEA), Reactome (reactome.org (35)), and the STRING database (36). GSEA analyses revealed that HOXB6 and/or HOXB8 regulated several cancer-related pathways such as cell cycle, apoptosis, and cellular senescence (Fig.4A, B). This supports the idea that HOXB6 and HOXB8 regulate these processes in PANC-1 cells.

**Figure 4.**
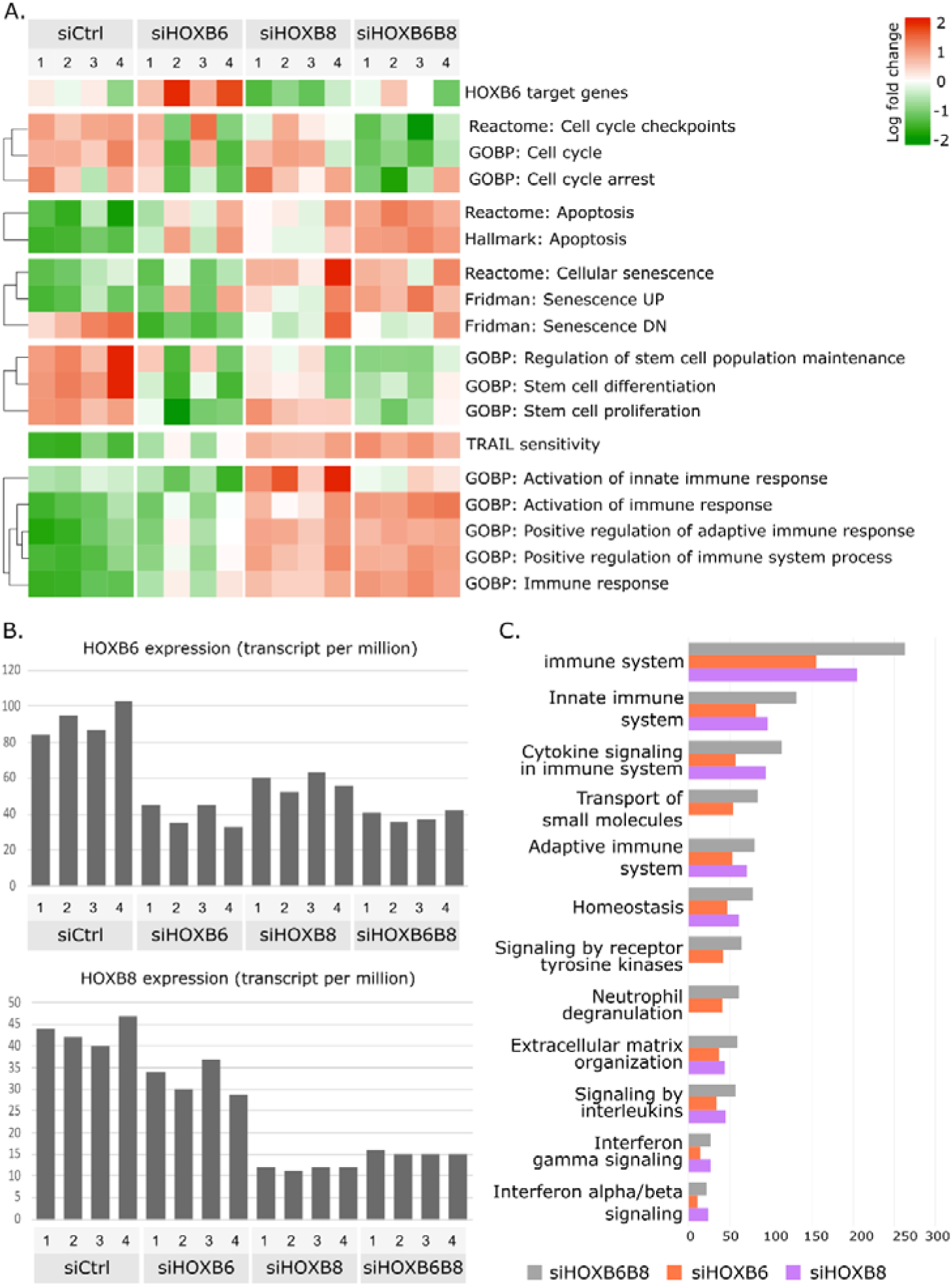
Transcription profiles and pathway analyses of siHOXB6, siHOXB8 and siHOXB6B8 in PDAC. (A) Gene set enrichment analyses, N=4 per condition. (B) Transcript per million of HOXB6 and HOXB8 genes. A disparity in the results between siHOXB6 replicates is observed (A, samples 1/3 vs 2/4) and is correlated with KD efficiency. (C) Number of genes identified in Reactome enrichment pathways up-regulated in siHOXB6, siHOXB8 and siHOXB6B8.

Interestingly, siHOXB6 and siHOXB8 influenced gene expression differently. Loss of HOXB6 predominantly led to enhanced expression of known HOXB6 target genes, while siHOXB8 reduced expression of these same genes. The effect of HOXB6 or HOXB8 deficiency was context dependent, as effects on gene expression varied depending on the biological processes investigated. For example, siHOXB6 reduced expression of genes associated with the activation of innate immune response, while siHOXB8 resulted in increased expression of genes in the same pathway. The phenotype of siHOXB6B8 PANC-1 cells had an intermediate effect suggesting that HOXB6 and HOXB8 act on the same genes to modulate gene expression. Down-regulation of genes involved in cell cycle and stem cell proliferation were mainly observed in siHOXB6 and double KDs (Fig.4A). However, we also observed cumulative effects of siHOXB6B8 compared to single KDs. This resulted in higher expression of gene sets for apoptosis, senescence, TRAIL (TNF-related apoptosis-inducing ligand) sensitivity, and activation of immune system in siHOXB6B8. STRING functional enrichment analyses using Reactome pathways (Fig.4C) confirmed GSEA observations and showed an over-representation of genes regulating the immune system in the KD cells, suggesting that HOX genes promote tumor cell survival through regulating immune evasiveness.

To identify which pathways were directly regulated by HOXB8, siHOXB8 differentially expressed gene list was compared with ChIP-seq data from HOXB8 chromatin immunoprecipitation (ChIP) in PANC-1 cells (37). HOXB8 promoter occupancy was observed in only a few genes with reduced siHOXB8 expression. These genes include stem cell markers CD24 and LGR5, pancreatic progenitor genes PDX1 and NKX6-2, and HOX genes such as HOXB6 (Fig.S3A) suggesting that HOXB8 primarily acts as a transcriptional repressor in PDAC. STRING functional enrichment analyses showed that HOXB8 promoter occupancy was observed in immune system, ECM organization and death receptor signaling pathway genes. Expression of the same genes was up-regulated upon loss of HOXB8 (Fig.S3B). HOXB8 promoter occupancy in combination with enhanced expression in siHOXB8 RNAseq samples suggests that HOXB8 directly inhibits these genes.

### HOXB6 and HOXB8 modulate immune response

RNAseq of siHOXB6 and siHOXB8 PANC-1 cells indicated that HOX genes regulate the transcription of genes associated with cellular immune response and possibly tumor immune evasiveness. To assess if HOXB6 and HOXB8 affect tumor cell-immune interactions *in vivo* the correlation between HOX gene expression and estimated immune cell infiltration was assessed using TIMER algorithms (http://timer.cistrome.org/ (18)).

The TIMER immune association module showed that HOXB6 and HOXB8 expression was positively correlated with the presence of immunosuppressive cell infiltrates such as myeloid-derived suppressor cells (MDSC) and tumor-associated fibroblasts (Fig.5A, n=179) in PDAC samples, further supporting an association between HOX gene expression and immune response in PDAC.

**Figure 5.**
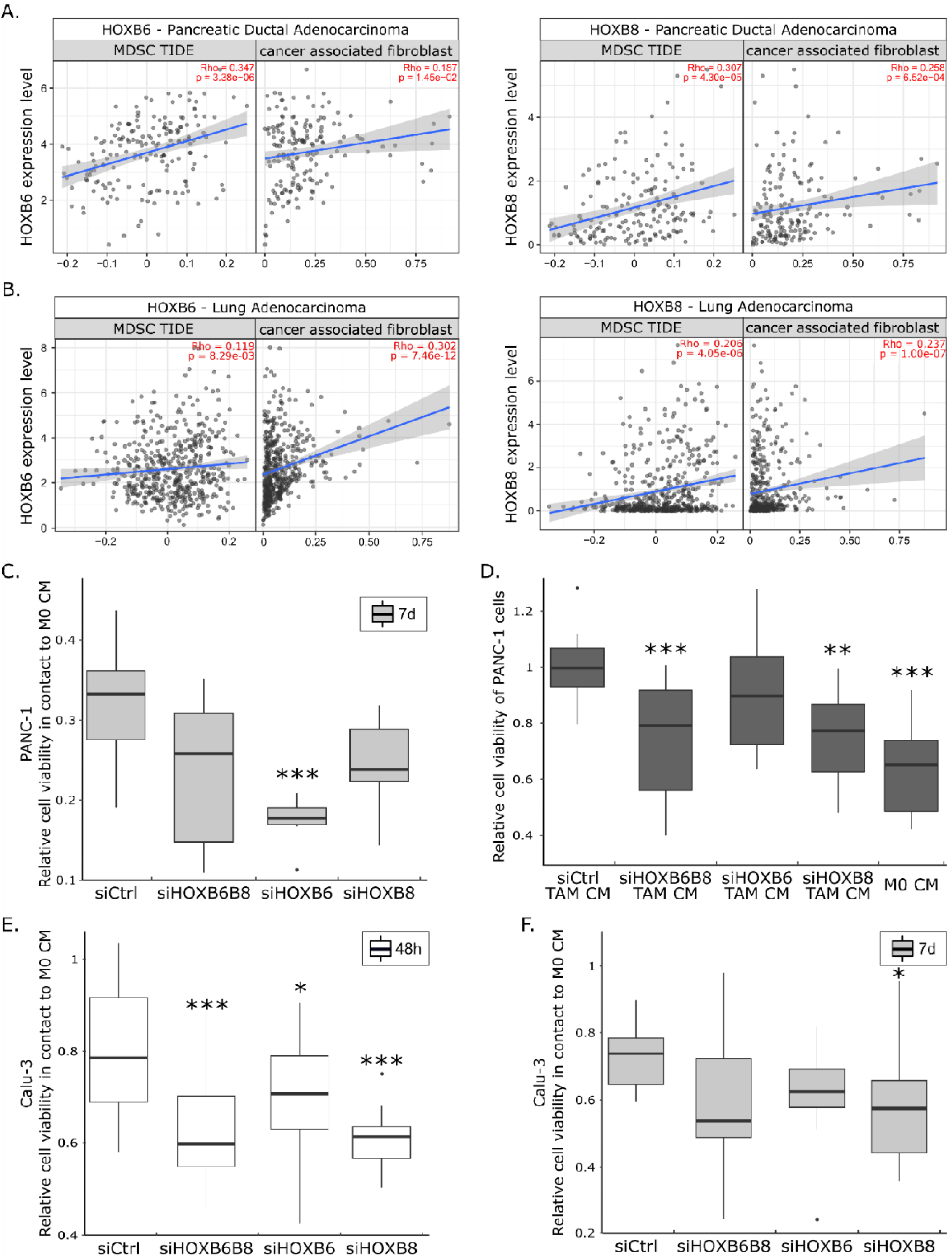
HOXB6 and HOXB8 modulate immune response. Correlation of HOXB6 and HOXB8 expression (log2 TPM) with the infiltration level of myeloid-derived suppressor cells (MDSC) and cancer associated fibroblasts estimated by TIMER in PDAC (A) and LUAD (B). Relative viability of PANC-1 cells (C, D) or Calu-3 cells (E, F) treated with macrophage CMs. MTT assays quantifying the relative cell viability of PANC-1 transfected cells (C, 7 days post transfection) or Calu-3 transfected cells (2 (E) or 7 (F) days post transfection) after treatment with M0 CM and (D) of PANC-1 cells exposed to conditioned mediums from siHOX-specific TAMs. Measurements were derived from three independent experiments. Tukey’s post-hoc test significances are indicated by stars compared to the control when significant. * p < 0.05 ** p < 0.01 *** p < 0.001. The box extends from the first to the third quartile (Q1/Q3) with a line representing the median. The whiskers extending from both ends of the box indicate variability outside Q1 and Q3. Everything outside is represented as an outlier.

To assess if HOXB6 and/or HOXB8 expression influenced immune cell-pancreatic tumor interactions, co-culture experiments of siHOXB6 and siHOXB8 PANC-1 cells with differentiating THP-1 monocytes were performed. THP-1 monocytes were stimulated with PMA to induce differentiation to M0 macrophages and conditioned medium (CM) was collected. The cellular response of siHOXB6 and siHOXB8 PANC-1 cells to M0 CM (see Fig.S4 for an overview of the experimental design) was assessed. As expected, exposure to M0 CM decreased cell number in siCtrl cells, HOX KD cell numbers decreased even further with siHOXB6 cells being the most sensitive (-45% compared to siCtrl+M0) to anti-tumorigenic proprieties of M0 macrophages (Fig.5C).

Cancer cells promote the differentiation of M0 macrophages into tumor-associated macrophages (TAM) which produce pro-oncogenic signals and promote tumor cell survival (38, 39). To test if HOXB6 and/or HOXB8 contribute to the differentiation of M0 macrophages to TAM, M0 macrophages were co-cultured with control and HOX KD cells (Fig.S4B), the CM of differentiated macrophages was collected, PANC-1 cells treated with CM, and cell viability assessed. CM from control cells promoted TAM differentiation and subsequently PANC-1 cell viability (Fig.5D). In contrast, CM from siHOXB8 and siHOXB6B8-differentiated macrophages significantly decrease cell viability (-24 and -26%, respectively (Fig.5D)) showing that M0 macrophages co-cultured with HOX KD cells maintained their capability to promote tumor cell death.

To determine if the role of HOXB6 and/or HOXB8 in regulating the immune response is PDAC specific or a general mode of action of HOX genes, we performed a TIMER association analysis of HOXB6 and HOXB8 expression in all available TCGA cancer samples. From the 40 cancer types studied, HOXB6 (Fig.S5) and HOXB8 (Fig.S6) expression correlated with cancer associated fibroblast and myeloid-derived suppressor cell infiltration in 7 different cancer types. From this list, only the lung carcinoma (LUAD) had a similar correlation pattern between HOXB8 expression and monocyte and macrophage infiltration (Fig.5B, S6). Therefore, we studied if HOX genes controlled macrophage-cancer cell interactions in Calu-3 cells, a LUAD cell line.

To assess if HOXB6 and/or HOXB8 expression influenced immune cell-tumor interactions in lung adenocarcinoma, we tested the sensitivity of siHOXB6 and/or siHOXB8 Calu-3 cells to exposure with M0 CM (Fig.S4C). As observed for PANC-1, exposure to M0 CM decreased cell number and HOX KDs cells were more sensitive to the anti-tumorigenic proprieties of M0 macrophages (-20%, -15%, and -21% compared to siCtrl+M0 for 7-day siHOXB6B8, siHOXB6, and siHOXB8, respectively) (Fig.5F). However, high variations were observed between replicates due to variable KD efficiencies 7 days post transfection (Fig.S7). KD efficiency was more consistent 48 hours post transfection (Fig.S7), thus sensitivity of KD cells to M0 CM in 48h transfected Calu-3 (Fig.5E) was assessed. Cell viability of HOX KD Calu3 cells was reduced upon M0 CM exposure (-21%, -14%, and -24% compared to siCtrl+M0 for 2-day siHOXB6B8, siHOXB6, and siHOXB8, respectively). We also assessed if Calu-3 cells contributed to the differentiation of M0 macrophages to TAM. Our results showed that M0 macrophages were losing their anti-tumorigenic proprieties upon incubation with Calu-3 cells and that the collected medium did not affect Calu-3 viability (Fig.S7), suggesting that Calu-3 cells were not responding to Calu-3 TAM CM.

Our data show that HOX genes promote PDAC cell-induced differentiation of TAM, as well as PDAC and LUAD cell resistance to M0 macrophages suggesting that elevated HOXB6 and HOXB8 expression levels contribute to an immune microenvironment that promotes enhanced tumor cell survival in different cancer types at least *in vitro*.

## DISCUSSION

Hox homeobox genes are well characterized transcription factors that establish the anterior-posterior axis in vertebrates and are critical for the morphogenesis of multiple organs (40). Gene duplication has resulted in the presence of several different HOX clusters (A to D) with paralog groups expressed in overlapping domains having similar DNA binding abilities. In addition to their roles in development, aberrant expression has also been associated with tumor formation and HOX genes regulate tumor cell proliferation, growth, migration and apoptosis (12). In these instances both loss and gain of HOX gene expression may promote tumorigenesis and the same HOX gene can act as tumor suppressor or proto-oncogene depending on the cellular context (41). In PDAC, aberrant expression of several HOX transcriptions factors has been described and functional studies have shown that HoxA10 (25), HoxB7 (24) and HoxC11 (42) promote pancreatic cancer development while HoxA1 (43), HOXB1 (44) and HoxB3 (23) may act as tumor-suppressors. However, the roles of HOXB6 and HOXB8 in PDAC and how these factors may interact has not been studied extensively. HOXB6 expression has been reported previously in mouse pancreatic fibroblasts which were not associated with the tumor stroma (28). Our expression analysis showed that, although abundant expression was observed in embryonic mesenchymal cells, HOXB6 was predominantly expressed in malignant ductal cells in PDAC while HOXB8 expression was limited to few cells during pancreas development and expression was virtually absent in adult pancreas but initiated in PDAC ductal cells suggesting that upregulation of these genes contribute to tumor formation.

Our gene expression analysis of HOXB6 and HOXB8 KD cells suggested that HOXB6 and HOXB8 promote PDAC tumorigenesis, but despite of having high molecular similarities the cellular mechanisms involved differ: HOXB6 mainly promotes expression of genes important for stem cell differentiation whereas HOXB8 appears to play a key function in immune response regulation. Interestingly, loss of HOXB6 and HOXB8 reduced stem cell proliferation and induced an immune response to tumor cells suggesting that HOXB6 and HOXB8 act in parallel (Fig.5).

Single cell RNAseq analysis showed that HOXB6 expression is present in malignant ductal cells, a cell population that also includes PDAC stem cells. Interestingly, a previous transcriptional characterization of PDAC stem cells showed that these cells have abundant HOXB6 expression suggesting that HOXB6 function promotes tumor stem cells (45). Moreover, our analysis of published HOXB8 ChIPseq data showed that HOXB8 binds to CD24 and LGR5 promoters in PANC-1 cells further supporting a function of these factors in tumor stem cells (37). Most importantly, we show that HOXB6 and HOXB8 KD impaired tumor cell proliferation, differentiation, and maintenance further supporting that HOXB6 and HOXB8 are important regulators of pancreatic cancer stem cells. HOXB6 expression correlated with BMI1 and SOX9 and previous studies have shown that SOX9 enhances tumor progression through BMI1 binding and p21 down-regulation which induces cell proliferation and evasion of apoptosis and senescence (46). These findings are similar to our observation where we observed a down-regulation of BMI1 in siHOXB6 and siHOXB6B8 cells which correlates with the induction of senescence, up-regulation of the tumor suppressor CDKN1A (p21^cip1/waf1^) and repression of tumor growth.

A categorization of genes highly expressed in PDAC and embryonic pancreas, showed that a large portion of genes (including HOXB6) were associated with the pancreatic progenitor subtype of PDAC, while HOXB8 expression was associated with pancreatic progenitor and immunogenic subtypes. The existence of an immunogenic PDAC has been discussed, and it has been suggested that the immune cell expression signature in this subtype is due to contamination by infiltrating B and T cells, while a gene expression pattern similar to pancreatic progenitor-like cells is also present (47). We observed impaired cell proliferation and colony formation in siHOXB6 PANC-1 cells which is in line with elevated HOXB6 expression in the pancreatic progenitor PDAC subtype. In contrast siHOXB8 PANC-1 cells exhibited impaired cell proliferation, but also suppressed induction of tumor-associated macrophages illustrating that HOXB8 contributes to both the pancreatic progenitor and immunogenic features observed in the immunogenic PDAC subtype.

siHOXB8 cells had enhanced expression of death receptor genes which control sensitivity of tumor cells to TRAIL, a cytokine produced in peripheral tissue, to promote tumor cell death (48). Specifically, increased expression of the trail receptor genes TNFRSF10B and TNFRSF10A was observed in siHOXB6 and siHOXB8 cells. Previous studies have shown that low levels of TRAIL receptors contribute to worse prognosis, while the presence of cytoplasmic TRAIL-R1 is a positive prognostic marker for PDAC patients (49, 50) and a combined treatment of gemcitabine with mesenchymal stem cells expressing TRAIL shows promising results in treating PDAC (51). Our findings suggest that HOXB6 and HOXB8 may modulate TRAIL resistance of PDAC with high HOX gene expressing tumors being less sensitive to treatments involving TRAIL and gemcitabine.

M0 conditioned medium affected cell viability of siHOXB6 PANC-1 and siHOXB6/8 Calu-3 cells and co-culture experiments of siHOXB8 cells with differentiating macrophages showed that HOXB8 promoted differentiation of tumor-associated macrophages. These results confirm that the changes in immune pathway gene expression observed in HOXB8 PANC-1 RNAseq data affect immune and cancer cell interactions (at least) *in vitro*. Correlation analysis between HOX gene expression and immune cell infiltration in human PDAC and LUAD tissues show that these mechanisms are also present *in vivo* in at least two different adenocarcinoma types that had relatively high HOXB8 expression and co-culture experiments performed in LUAD and PANC-1 showed that the regulation of immune-cancer cell interaction of HOX genes is conserved. Moreover, a recent publication showed that similar HOX genes control similar processes in endometrial cancer with a HOX expression score being proposed to predict among other parameters the efficacy of immunotherapy (52). This is significant as it suggests that HOXB6 and HOXB8 expression (in at least pancreatic and lung adenocarcinoma) enhance immunosuppressive effects of cancer cells by promoting immune evasiveness of macrophages and thereby contributing to an immunosuppressive microenvironment affecting macrophage differentiation.

Our findings indicate that HOX genes play an important role in regulating cell proliferation, cell death, immune response, and treatment resistance in pancreatic cancer cells. These results are consistent with the pathways correlating with HOXB6 and HOXB8 in PDAC tissues and the immune cell infiltrates observed in PDAC and LUAD. Our data suggest that HOXB6 and HOXB8 regulate numerous oncogenic pathways to promote adenocarcinoma tumorigenesis.

## Ethics approval and consent to participate

Ethical permission has been obtained from the regional ethics committee in Lund (Dnr2012/593, Dnr 2015/241, Dnr 2018-579). The Surgical Ethics Committee of HUH approved the study protocol (Dnr. HUH 226/E6/06, extension TKM02 §66 17.4.2013).

## Availability of data and materials

The sequencing data related to fetal samples generated for this study can be found in the LUDC repository (www.ludc.lu.se/resources/repository) under the following accession numbers and are available upon reasonable request: LUDC2021.10.12 (bulk RNA sequencing data from fetal pancreata), and LUDC2021.10.18 (single cell RNA sequencing data from fetal pancreas). The PANC-1 sequencing data have been deposited with links to BioProject accession number PRJNA1043644 in the NCBI BioProject database (https://www.ncbi.nlm.nih.gov/bioproject/).

## Competing interests

The authors declare that they have no competing interests.

## Funding

Our research was funded by the Fredrik and Ingrid Thuring foundation (2020-00596), Swedish Research council (2020-0146), Novonordisk Foundation (NNF20OC0063485), the Swedish Cancer foundation (2022), the Royal Physiographic Society (2022), the Lars Hiertas Minne foundation (2022) and the Cancer and Allergy Foundation (2022, 2023).

## Authors’ contributions

Conceptualization, LBB, RBP and IA; methodology, LBB, SB, KA, TS, JA, RBP, TK, JH, CH, HS; formal analysis, LBB; investigation, LBB, IA, TK, JH, CH, HS; data curation, KA, RBP, JA; writing original draft preparation, LBB; writing review and editing, IA; visualization, LBB, KA, RBP, JA, TK; supervision, IA; funding acquisition, LBB and IA. All authors have read and agreed to the published version of the manuscript.

## Acknowledgements

We thank Philip Kaldis, Lund University and Ulrike Nuber, TU Darmstadt for their comments on the manuscript. We thank Darcy Wagner, Lund University for the Calu-3 cell gift.

## Notes

The authors declare no potential conflicts of interest

### Competing Interest Statement

The authors have declared no competing interest.

