## Supplementary legends for "HOXB6 and HOXB8 control immune-cancer cell interactions in pancreatic cancer"

**Figure S1.** **Disease specific survival analyses in PDAC.** Kaplan–Meier survival curves for PDACs according to HOXB6 (A) or HOXB8 (B) protein expression. Kaplan–Meier analysis revealed that low HOXB6 expression associated to poorer OS compared to moderate expression (p=0.005) or high expression (p=0.034). No difference is seen between moderate and high HOXB6 expression. 5-year OS for patients with a low HOXB6 expression was 12.3% (95% CI 1.8-22.19%) compared to moderate 26.3% (95% CI 16.1-36.5%) and high expression 15.5% (95% CI 4.1-26.9%). Low HOXB8 expression is associated with poorer OS compared to moderate expression (p=0.007) or high expression (p=0.007). No difference in OS is seen between moderate and high HOXB8 expression. 5-year OS for patients with a low HOXB8 expression was 9.7% (95% CI 2.8-16.6%) compared to moderate 21.4% (95% CI 12.0-30.8%) and high expression 24.0% (95% CI 12.7-36.7%).

**Figure S2.** **Transfection efficiency determined by RT-qPCR analyses.** HOXB6 (A) and HOXB8 (B) knockdown efficiency 48h and 7 days after transfection. Expression values were calculated applying the −2∆∆CT algorithm. Estimated relative quantities were normalized for the expression value of the endogenous genes S18 and TBP and calibrated to the negative control samples. Measurements were derived from three independent experiments. Error bars indicate the standard deviation. Tukey’s post-hoc test significances calculated on the ∆CT values are indicated by stars compared to the control when significant. *** p < 0.001.

**Figure S3. HOXB8 ChiP-seq analyses.** (A)  Protein interaction (STRING) from the genes downregulated in siHOXB8 with a direct HOXB8 binding site identified by ChiP-seq. Lines represent the protein-protein association and the thickness indicates the strength of data support. The thinnest lines indicate a medium edge confidence and the thicker ones a high edge confidence. (B) Reactome pathways of genes upregulated in siHOXB8 with a direct HOXB8 binding site identified by ChiP-seq.

**Figure S4. Experimental set-up to quantify** **cell viability**. (A) Preparation of naïve macrophages (M0) conditioned medium (CM) and viability test on PANC-1 siHOX cells. (B) Preparation of CM from siHOX-specific tumor-associated macrophages (TAMs) and viability test on PANC-1 cells. (C) Preparation of naïve macrophages (M0) conditioned medium (CM) and viability test on Calu3 siHOX cells.

**Figure S5. Immune association with HOXB6 expression.** TIMER2.0 analyses of HOXB6 expression correlating with immune infiltration in all available TCGA cancer samples. Colored squares show significant results. Green rectangle shows cancer types where HOXB6 correlates with cancer associated fibroblasts, in pink, correlation with myeloid-derived suppressor cells (MDSC). Cancer abbreviations: ACC, Adrenocortical carcinoma; BLCA, Bladder Urothelial Carcinoma ; BRCA, Breast invasive carcinoma ; CESC, Cervical squamous cell carcinoma and endocervical adenocarcinoma ; CHOL, Cholangiocarcinoma ; COAD, Colon adenocarcinoma ; DLBC, Lymphoid Neoplasm Diffuse Large B-cell Lymphoma ; ESCA, Esophageal carcinoma ; GBM, Glioblastoma multiforme ; HNSC, Head and Neck squamous cell carcinoma ; KICH, Kidney Chromophobe; KIRC, Kidney renal clear cell carcinoma; KIRP, Kidney renal papillary cell carcinoma; LGG, Brain Lower Grade Glioma; LIHC, Liver hepatocellular carcinoma; LUAD, Lung adenocarcinoma; LUSC, Lung squamous cell carcinoma; MESO, Mesothelioma; OV, Ovarian serous cystadenocarcinoma; PAAD, Pancreatic adenocarcinoma; PCPG, Pheochromocytoma and Paraganglioma; PRAD, Prostate adenocarcinoma; READ, Rectum adenocarcinoma; SARC, Sarcoma; STAD, Stomach adenocarcinoma; SKCM, Skin Cutaneous Melanoma; TGCT, Testicular Germ Cell Tumors; THCA, Thyroid carcinoma ; THYM, Thymoma; UCEC, Uterine Corpus Endometrial Carcinoma; UCS, Uterine Carcinosarcoma; UVM, Uveal Melanoma.

**Figure S6. Immune association with HOXB8 expression.** TIMER2.0 analyses of HOXB8 expression correlating with immune infiltration in all available TCGA cancer samples. Colored squares show significant results. Green rectangle shows cancer types where HOXB8 correlates with cancer associated fibroblasts, in pink, correlation with myeloid-derived suppressor cells (MDSC). Cancer abbreviations: ACC, Adrenocortical carcinoma; BLCA, Bladder Urothelial Carcinoma ; BRCA, Breast invasive carcinoma ; CESC, Cervical squamous cell carcinoma and endocervical adenocarcinoma ; CHOL, Cholangiocarcinoma ; COAD, Colon adenocarcinoma ; DLBC, Lymphoid Neoplasm Diffuse Large B-cell Lymphoma ; ESCA, Esophageal carcinoma ; GBM, Glioblastoma multiforme ; HNSC, Head and Neck squamous cell carcinoma ; KICH, Kidney Chromophobe; KIRC, Kidney renal clear cell carcinoma; KIRP, Kidney renal papillary cell carcinoma; LGG, Brain Lower Grade Glioma; LIHC, Liver hepatocellular carcinoma; LUAD, Lung adenocarcinoma; LUSC, Lung squamous cell carcinoma; MESO, Mesothelioma; OV, Ovarian serous cystadenocarcinoma; PAAD, Pancreatic adenocarcinoma; PCPG, Pheochromocytoma and Paraganglioma; PRAD, Prostate adenocarcinoma; READ, Rectum adenocarcinoma; SARC, Sarcoma; STAD, Stomach adenocarcinoma; SKCM, Skin Cutaneous Melanoma; TGCT, Testicular Germ Cell Tumors; THCA, Thyroid carcinoma ; THYM, Thymoma; UCEC, Uterine Corpus Endometrial Carcinoma; UCS, Uterine Carcinosarcoma; UVM, Uveal Melanoma.

**Figure S7.** **Transfection efficiency determined by RT-qPCR analyses and Calu-3 sensitivity to TAM.**  HOXB6 (A) and HOXB8 (B) knockdown efficiency 48h and 7 days after transfection in Calu-3 cells. Expression values were calculated applying the −2∆∆CT algorithm. Estimated relative quantities were normalized for the expression value of the endogenous genes S18 and TBP and calibrated to the negative control samples. Measurements were derived from three independent experiments. Error bars indicate the standard deviation. Tukey’s post-hoc test significances calculated on the ∆CT values are indicated by stars compared to the control when significant. (C) MTT assay quantifying the relative cell viability of Calu-3 cells exposed to conditioned mediums from siHOX-specific TAMs, M0 or control culture medium. * p < 0.05, *** p < 0.001.

**Tables**

**Table S1.** Primer sequences for qPCR.

**Table S2.** Identification of genes controlling human pancreas development and subtype-specific pancreatic cancer

**Table S3.** Subtype-specific genes and the associated literature

**Table S4.** Overall survival analyses

**Table S5.** HOXB6 co-expression analyses in embryo tissues using Spearman's rank correlation method.

**Table S6.** HOXB6 co-expression analyses in normal adult tissues using Spearman's rank correlation method.

**Table S7.** HOXB6 co-expression analyses in PDAC using Spearman's rank correlation method.

**Table S8.** HOXB8 co-expression analyses in PDAC tissues using Spearman's rank correlation method.

**Table S9.** Pathways identified by Reactome using significant co-expressed genes with both HOXB6 and HOXB8 in PDAC.

**Table S10.** Pathways identified by Reactome using positively co-expressed genes with HOXB6 in PDAC.

**Table S11.** Pathways identified by Reactome using negatively co-expressed genes with HOXB6 in PDAC.

**Table S12.** Pathways identified by Reactome using positively co-expressed genes with HOXB8 in PDAC.

**Table S13.** Pathways identified by Reactome using positively co-expressed genes with HOXB6 in embryo.

**Table S14.** Pathways identified by Reactome using positively co-expressed genes with HOXB6 in adult pancreas

**Table S15.** Deregulated genes between siHOXB6 and control transcriptomes.

**Table S16.** Deregulated genes between siHOXB8 and control transcriptomes.

**Table S17.** Deregulated genes between siHOXB6B8 and control transcriptomes.
