## Supplementary material for "HOXB6 and HOXB8 control immune-cancer cell interactions in pancreatic cancer": TableS3

| **Gene** | **specificity** | **logFC** | **linked to PC** | **Functional studies in pancreatic cancer cells** |
| --- | --- | --- | --- | --- |
| **PLAC8** | 4 subtypes | >1.5 | Tatura et al., 2020  Kinsey et al., 2014  Kaistha et al., 2016 | Tatura et al., 2020  Kinsey et al., 2014  Kaistha et al., 2016 |
| **S100P** | 4 subtypes | >1.5 | Nakayama et al., 2019  Arumugam et al., 2005 | Nakayama et al., 2019  Arumugam et al., 2005 |
| **ZBED2** | 4 subtypes | >1.5 | Somerville et al., 2020 | Somerville et al., 2020 |
| **DHRS9** | 4 subtypes | <1.5 | Li et al., 2020  Wu et al., 2018 | Li et al., 2023 |
| **S100A2** | 4 subtypes | <1.5 | Ochuchida et al., 2007 | Li et al., 2021 |
| **IL1A** | 4 subtypes | <1.5 | Brunetto et al., 2019  Tjomsland et al., 2016 | Brunetto et al., 2019  Tjomsland et al., 2016 |
| **DLGAP5** | Squamous tumours | 4,4824 | Ke et al., 2020  Zhou et al., 2018 | [Mu-Jing Ke et al., 2020](https://pubmed.ncbi.nlm.nih.gov/?term=Ke+MJ&cauthor_id=32782440) |
| **PTTG1** | Squamous tumours | 4,3254 | Zhang et al., 2008  Lin et al., 2013  Long et al., 2016 | Long et al., 2023 |
| **SEMA3C** | Squamous tumours | 3,6704 | Shang 2019  Xu et al., 2017 | Zhang et al., 2022 |
| **CKS2** | Squamous tumours | 3,6419 | Li et al., 2018 | Li et al., 2018 |
| **GINS1** | Squamous tumours | 3,1229 | Azim et al., 2016  Zhou 2017 | Bu et al., 2023 |
| **CCNA2** | Squamous tumours | 3,0065 | Dong et al., 2019  Long et al., 2016  Zhou et al., 2018 | Jiang et al., 2020  Jiang et al., 2022 |
| **CYB5R2** | Squamous tumours | 2,81116 | Ming et al., 2015  Liu et al., 2015  Wong et al., 2020 |  |
| **DKK1** | Squamous tumours | 2,80433 | Wei et al., 2020  Kimura et al., 2019  Igbinigie et al., 2019  Sato et al., 2010 | Kimura et al., 2021 |
| **SHCBP1** | Squamous tumours | 2,76508 | Luo 2020 | Luo et al., 2020 |
| **CCNB1** | Squamous tumours | 2,76499 | Zhou et al., 2018  Zhang et al., 2018 | Zhang et al., 2018 |
| **KPNA2** | Squamous tumours | 2,3203 | Rachidi et al., 2013 | Zhou et al., 2021 |
| **CMTM3** | Squamous tumours | 2,26138 | Zhou et al., 2021 | Zhou et al., 2021 |
| **HRH1** | Squamous tumours | 1,93802 | Salmon et al., 2020 | Salmeron et al., 2021 |
| **ZNF532** | Squamous tumours | 1,74962 | Zhou et al., 2015 |  |
| **IL1RAP** | Squamous tumours | 1,73442 | Zhang et al., 2022 | Zhang et al., 2022 |
| **PDLIM7** | Squamous tumours | 1,50757 | Ramezankhani et al., 2022 |  |
| **AMIGO2** | Squamous tumours | 1,4428 | Shen et al., 2017 |  |
| **MAP4K4** | Squamous tumours | 1,42381 | Liang et al., 2015  Fu et al., 2018  Zhao et al., 2015  Chen et al., 2019 | Zhao et al., 2015 |
| **SLC7A7** | Squamous tumours | 0,79225 |  |  |
| **ARL4C** | Squamous tumours | 0,74632 | Harada et al., 2021  Chen et al., 2021 | Harada et al., 2021  Chen et al., 2021 |
| **KLF7** | Squamous tumours | 0,70081 | Gupta et al., 2020  Yu et al., 2019 | Gupta et al., 2020 |
| **TTK** | pancreatic progenitor tumours | 3,471 | Kaistha, 2014 | Lui, 2014  Yao et al., 2021 |
| **NUF2** | pancreatic progenitor tumours | 3,2968 | Hu et al., 2015 | Hu et al., 2015 |
| **RAD51AP1** | pancreatic progenitor tumours | 2,78345 | Liu et al., 2022 | Liu et al., 2022 |
| **HOXB6** | pancreatic progenitor tumours | 1,90783 | Segara et al., 2005  Garcia et al., 2020 | Garcia et al., 2020 |
| **CORO2A** | pancreatic progenitor tumours | 1,74244 | Tang et al., 2018  Xi et al., 2023 |  |
| **HOXB3** | pancreatic progenitor tumours | 1,72247 | Yang et al., 2016 | Yang et al., 2016 |
| **SLC16A5** | pancreatic progenitor tumours | 0,89256 | Yu et al., 2020 |  |
| **TMC7** | pancreatic progenitor tumours | 0,52086 | Cheng et al., 2019 | Cheng et al., 2015 |
| **E2F8** | immunogenic tumors | 3,9818 | Luo et al., 2021 |  |
| **MALL** | immunogenic tumors | 0,79755 | Liu et al., 2018 |  |
| **ABLIM3** | immunogenic tumors | 0,53176 |  |  |
| **TOP2A** | ADEX tumours | 5,1226 | Zhou 2017  Pei, 2017 |  |
| **CENPF** | ADEX tumours | 4,0768 | Cheng 2019  Zhou et al., 2018 | Cheng et al., 2019  Chen et al., 2021 |
| **MELK** | ADEX tumours | 3,6981 | Li et al., 2020  Chung et al., 2012 | Li et al., 2020 |
| **NMU** | ADEX tumours | 3,1305 | Gu et al., 2019 | Yoo et al., 2019 |
| **SHISA2** | ADEX tumours | 3,1096 | Zhu et al., 2017 | Chu et al., 2020 |
| **MMP11** | ADEX tumours | 2,6609 | Jungwhoi et al., 2019 Zhang et al., 2020 | Jungwhoi et al., 2019  Zhang et al., 2020 |
| **DPYSL3** | ADEX tumours | 2,58388 | Hiroshima et al., 2012  Wu et al., 2018  Kawahara et al., 2013 | Hiroshima et al., 2012  Kawahara et al., 2013 |
| **CRIP1** | ADEX tumours | 2,49753 | Zhang et al., 2018  Hao et al., 2008  Baumhoer et al., 2011 | Zhang et al., 2018 |
| **CDH11** | ADEX tumours | 2,47585 | Birtolo et al., 2017 | Birtolo et al., 2017 |
| **ANO1** | ADEX tumours | 2,21992 | Ardelenau et al., 2009 Bergmann et al., 2011 Sauter et al., 2014 | Sauter et al., 2015  Zhang et al., 2023 |
| **ANTXR1** | ADEX tumours | 2,15449 | Alcala et al., 2019 | Alcala et al., 2019 |
| **EPSTI1** | ADEX tumours | 1,88833 | Hastie et al., 2016  Ke, 2022 |  |
| **STIL** | ADEX tumours | 1,56067 | Ito et al., 2020 | Ito et al., 2020 |
| **ECT2** | ADEX tumours | 1,55328 | Zhang et al., 2008  Liu et al., 2018  Samuel et al., 2013 | Liu et al., 2018 |
| **S100A16** | ADEX tumours | 1,40267 | Tomiyama et al., 2018 | Tomiyama et al., 2018 |
| **RHBDL2** | ADEX tumours | 1,24143 | Khalid et al., 2019 | Chen et al., 2022 |
| **LY6E** | ADEX tumours | 1,16372 | Luo et al., 2016  Gou et al., 2007  Russ et al., 2021 |  |
| **SGIP1** | ADEX tumours | 0,7627 | Zhou et al., 2016 |  |
| **CAP1** | ADEX tumours | 0,6204 | Wu et al., 2019  Yamazaki et al., 2009 | Wu et al., 2019 |
| **OAS1** | ADEX tumours | 0,53655 | Wang, 2019  Lu et al., 2022 |  |
| **OAS3** | ADEX tumours | 0,52681 | Wang, 2019  Gao et al., 2022 |  |
