## Supplementary figures and images for "HOXB6 and HOXB8 control immune-cancer cell interactions in pancreatic cancer"

### Fig.S1

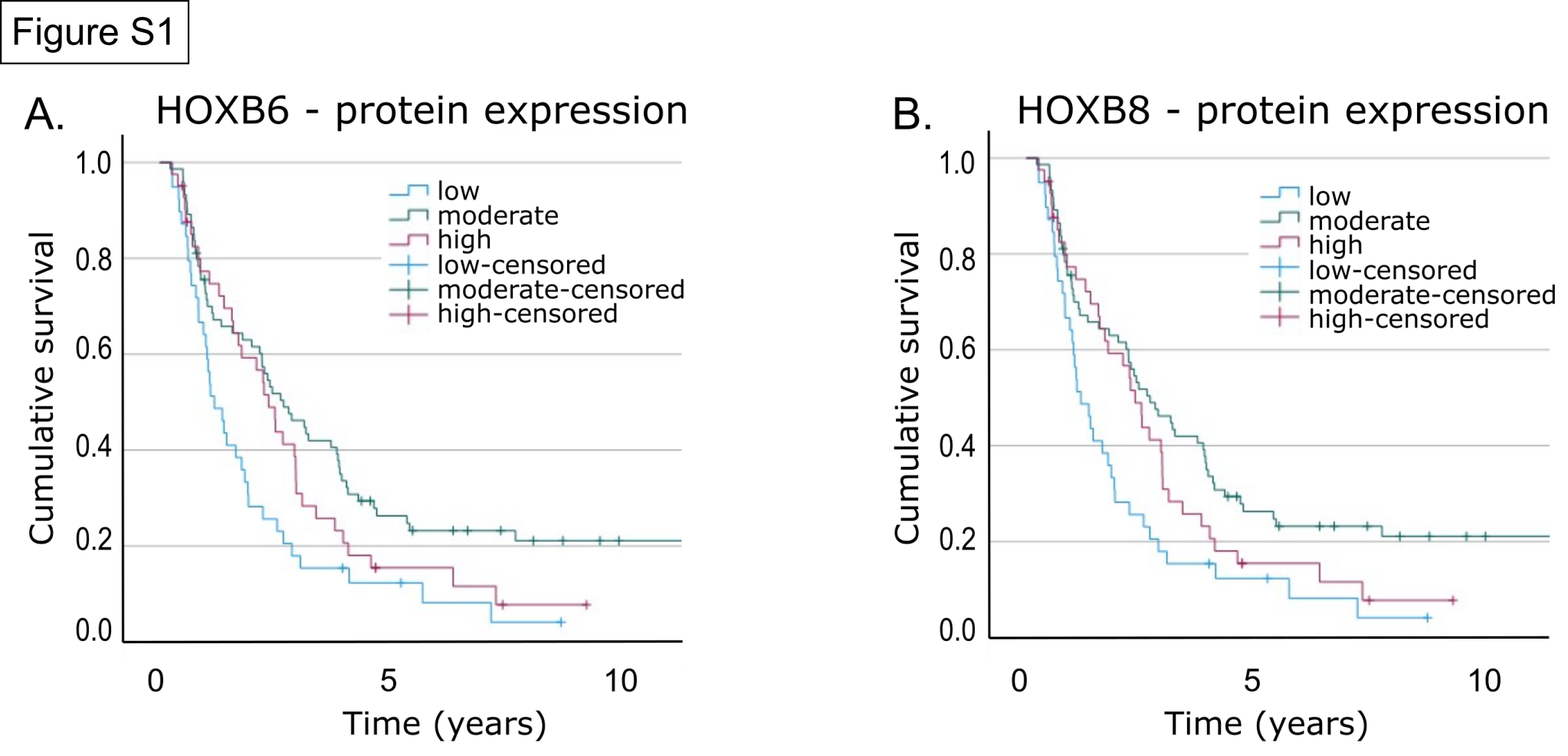

### Fig.S2

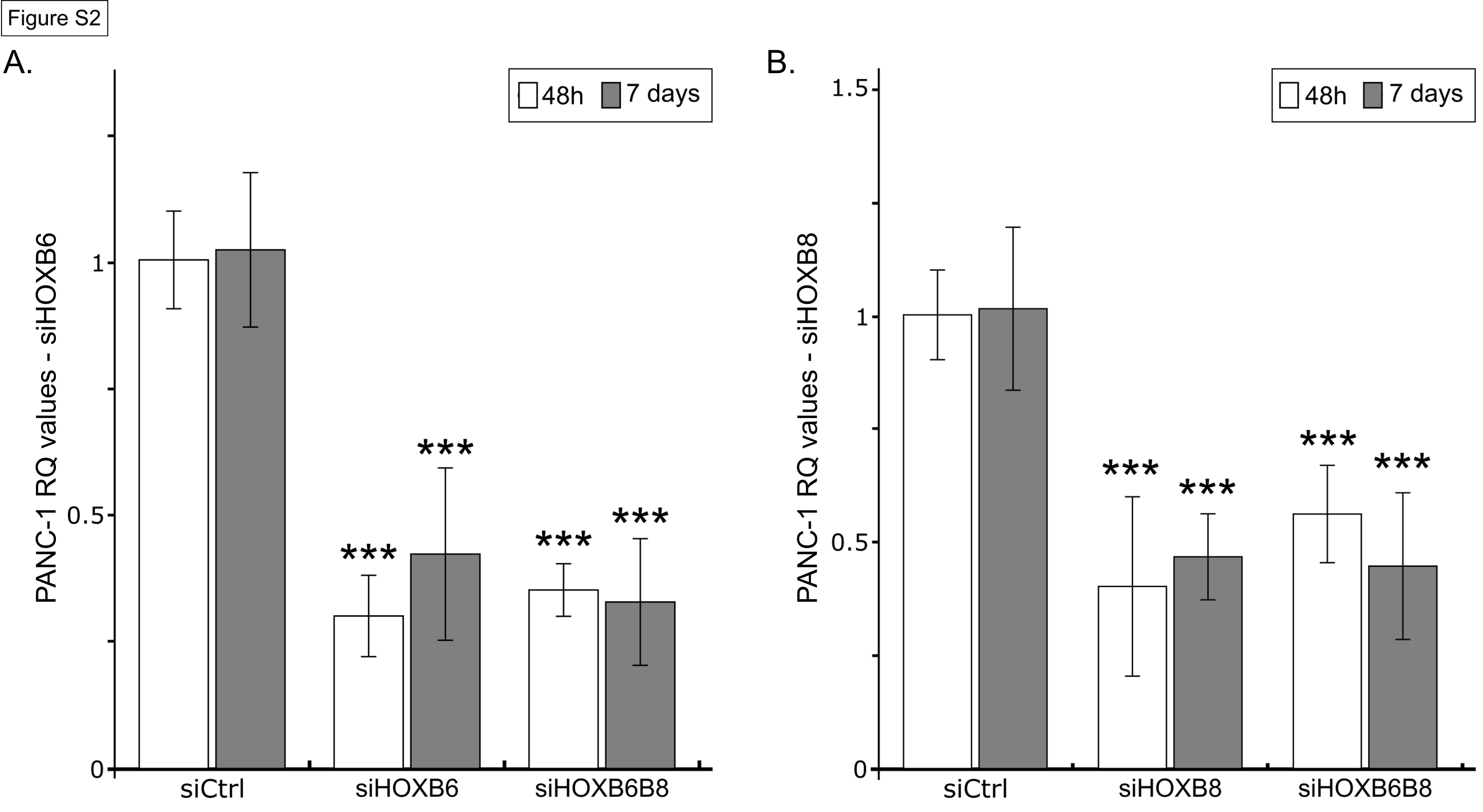

### Fig.S3

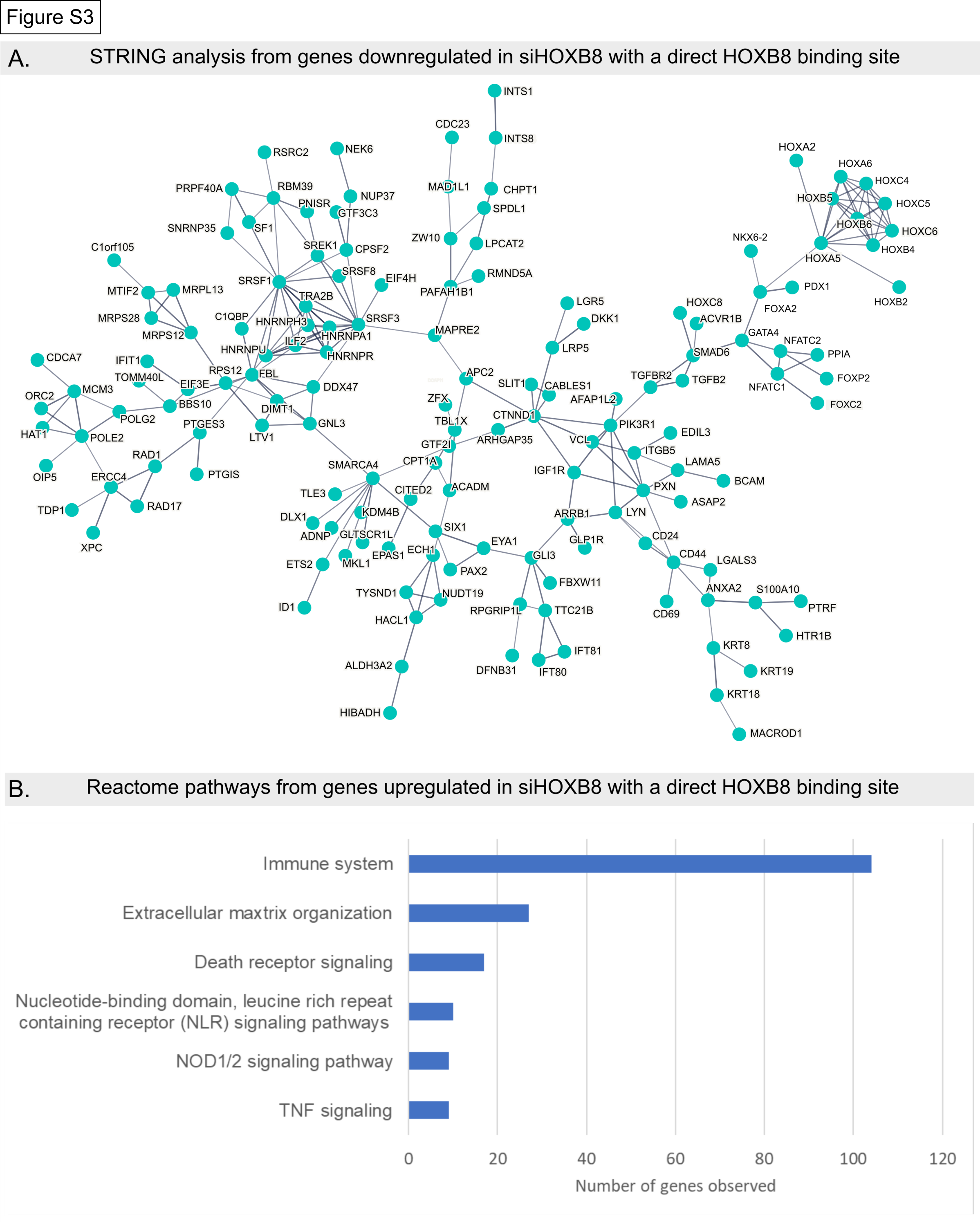

### Fig.S4

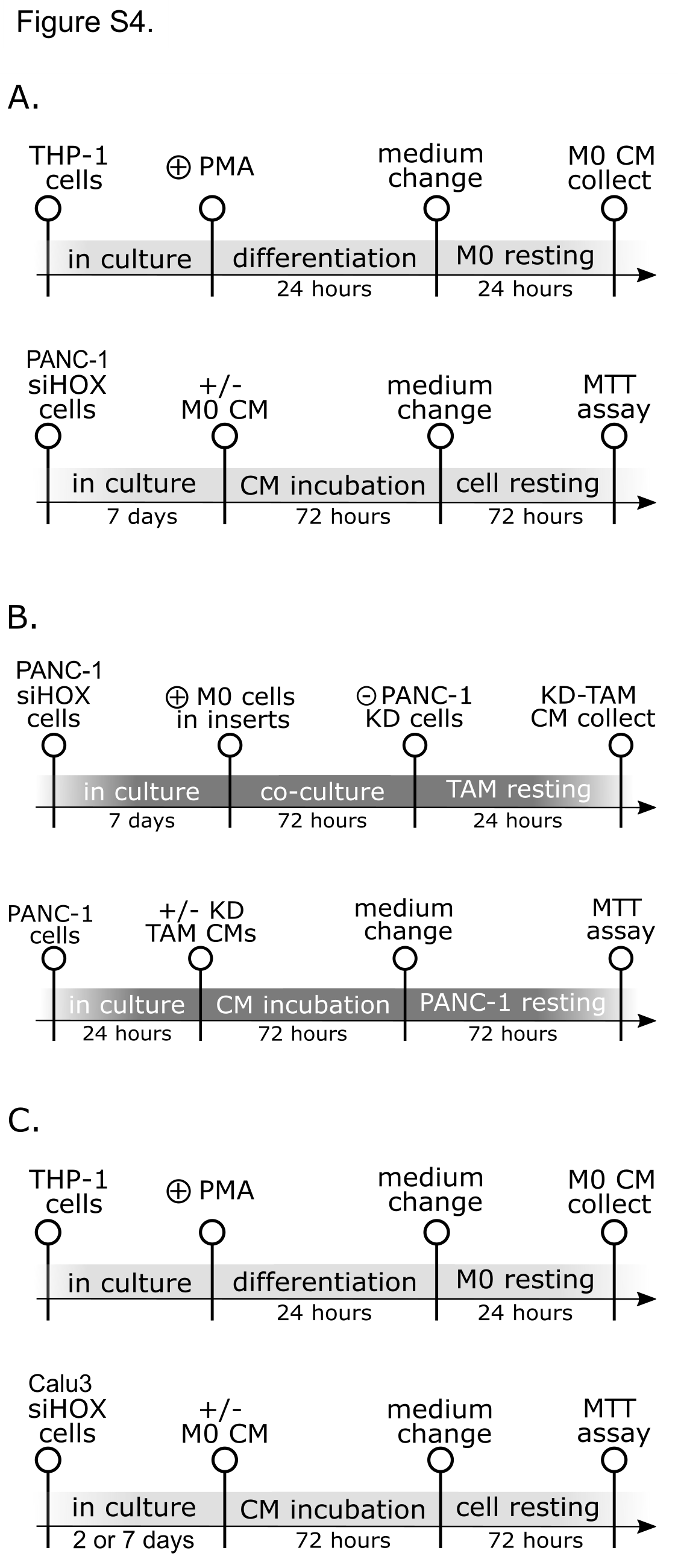

### Fig.S5

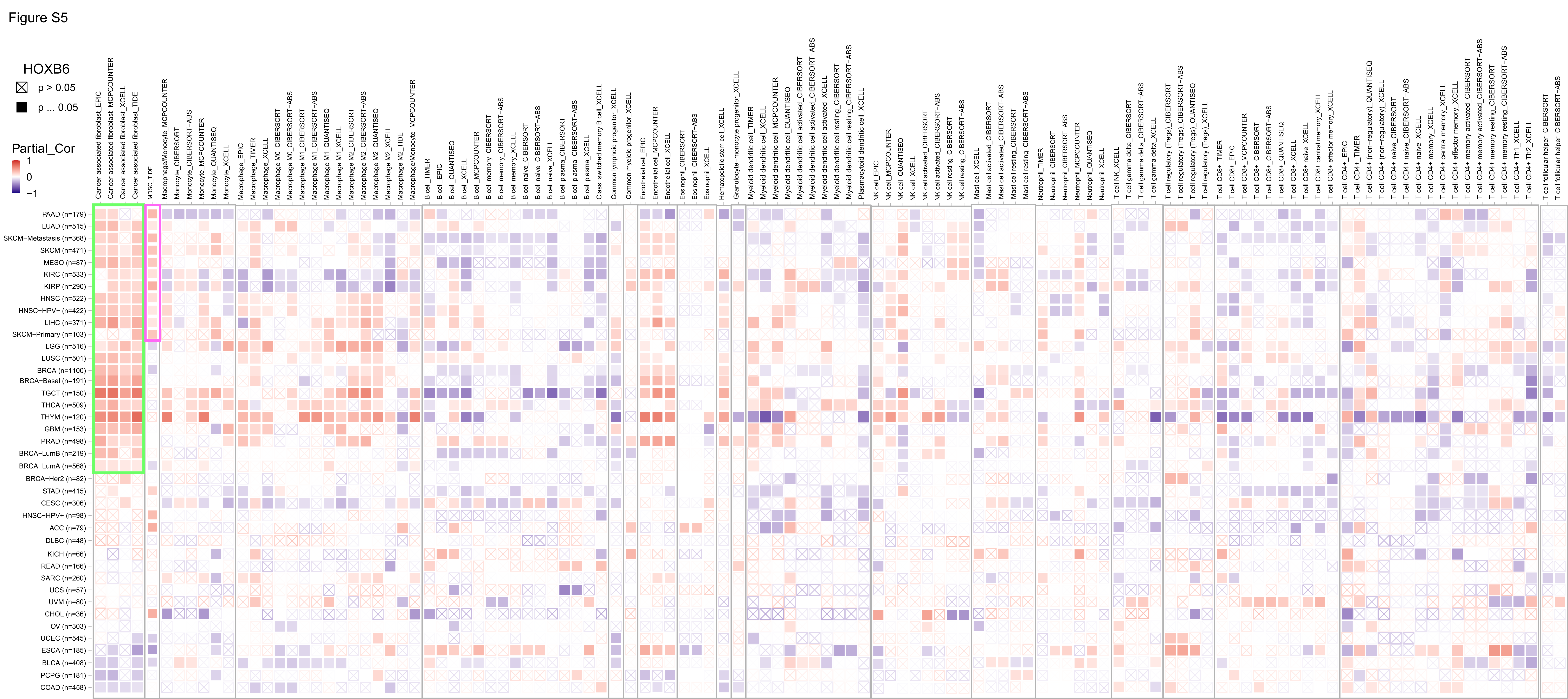

### Fig.S6

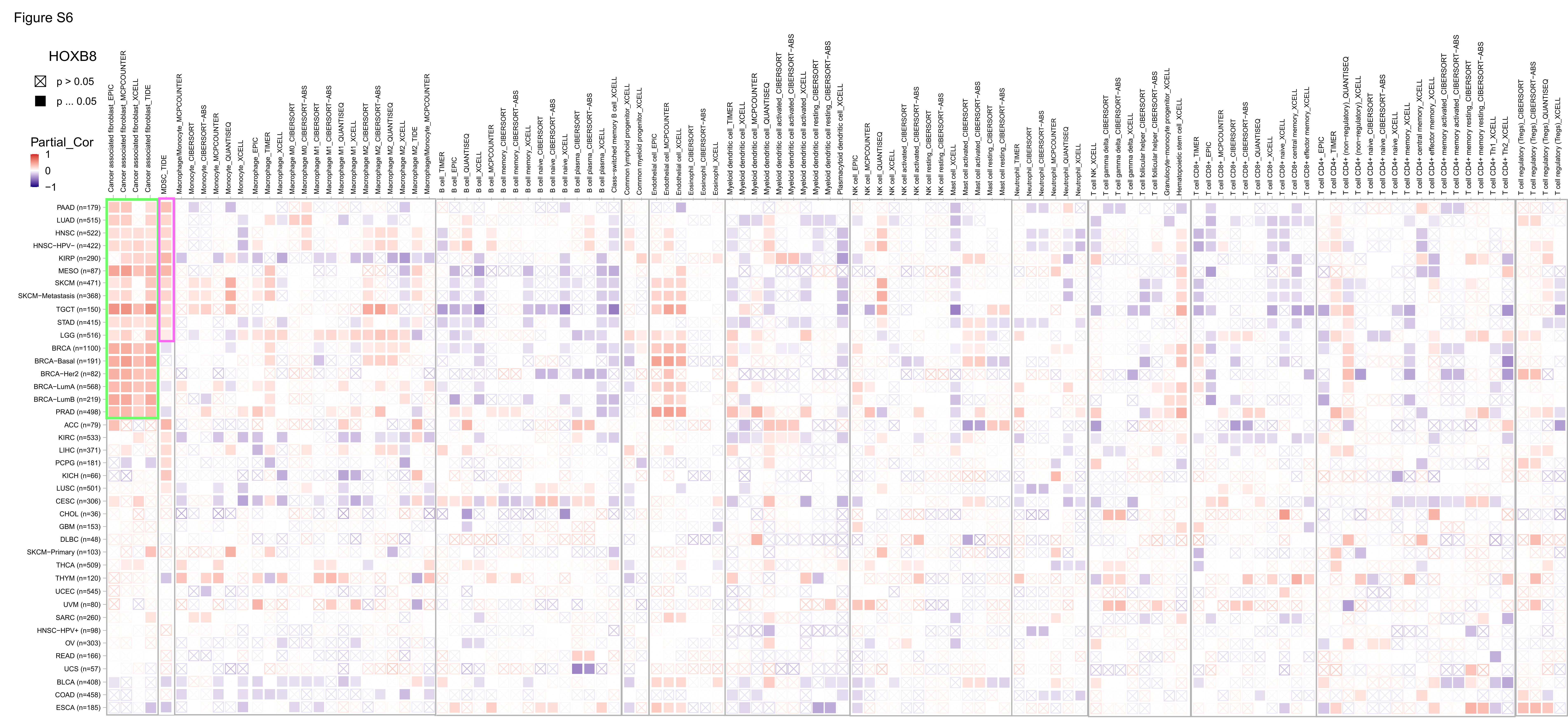

### Fig.S7

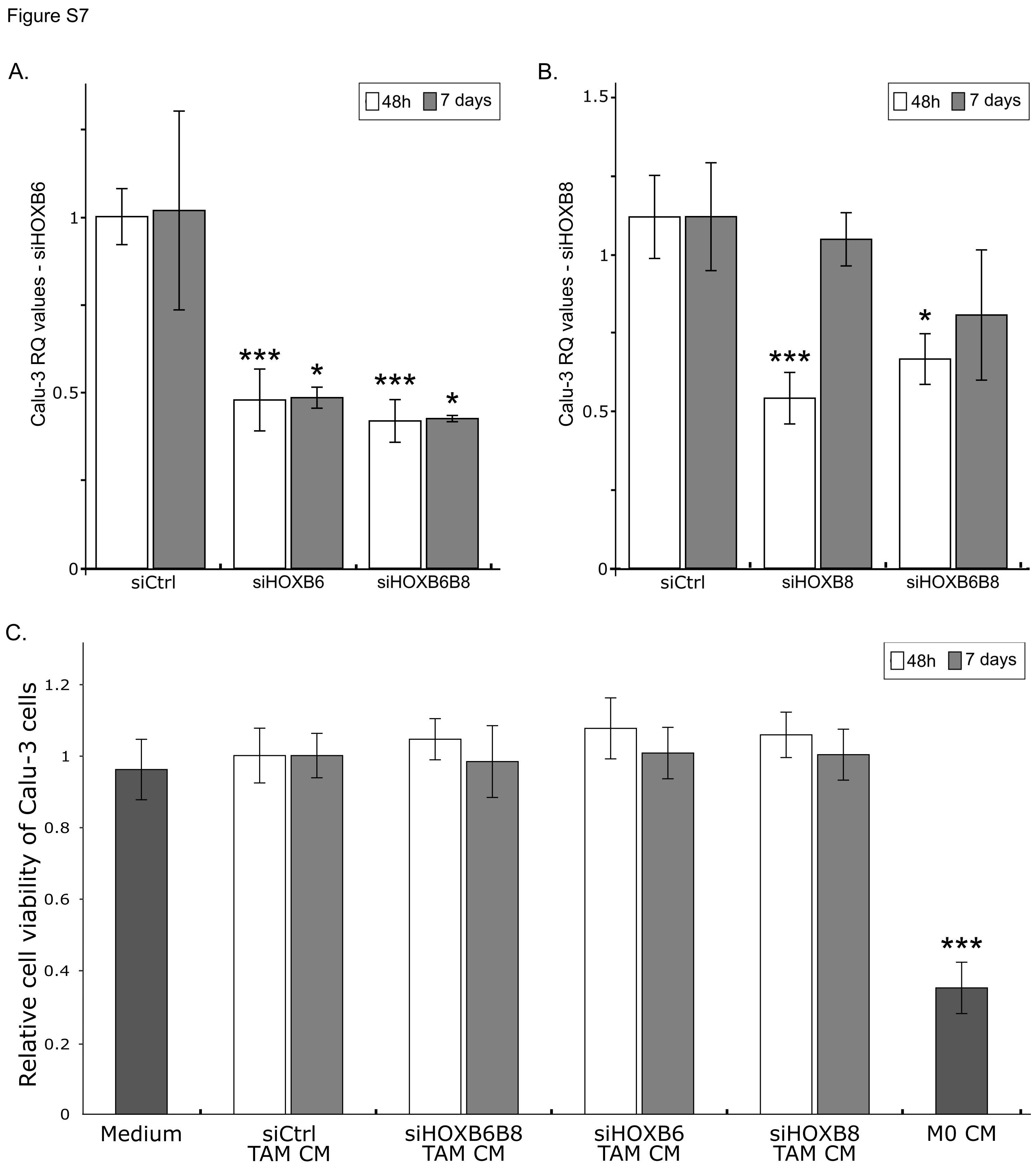
